# Population genomics of *Actinobacillus pleuropneumoniae* reveals lineage-associated K:O antigen and Apx toxin diversity

**DOI:** 10.64898/2026.10.19.759954

**Authors:** Jiamao Yang, Maria Hakimi, Xiao Hu, Maria J. Clavijo, Marcelo N. Almeida, Nubia R. Macedo, Huigang Shen, Ganwu Li

## Abstract

*Actinobacillus pleuropneumoniae* (APP) is the causative agent of porcine pleuropneumonia, causing high mortality and morbidity and substantial economic losses in the swine industry. Nineteen capsular (K) serotypes, defined by variation in capsular polysaccharide (CPS) biosynthesis loci, ten O serotypes, defined by variation in lipopolysaccharide (LPS) biosynthesis loci, characteristic Apx toxin profiles, and genomic diversity have been increasingly described; however, relationships among genomic lineage, surface antigen genotype, and Apx toxin repertoire remain incompletely resolved. Here, we analyzed 328 APP genomes, including 153 newly sequenced isolates and 175 public genomes from 21 countries. Core-genome SNP phylogeny and FastBAPS identified 23 genomic lineages. Nineteen capsular K types and 10 O-antigen types formed 26 K:O genotypes, of which 24 were restricted to a single lineage. Adjusted Wallace analysis showed greater concordance of K:O genotype with lineage than K type alone (0.881 versus 0.759), and with ApxI–ApxIII genotype (0.949 versus 0.787). ApxIVA-related sequence was detected in all genomes, while 56 complete loci revealed recurrent length variation within the N-terminus or the RTX repeat-rich region and an additional divergent ApxIVA-related homolog. In conclusion, our comprehensive population genomic study gives a more informative picture of APP genomics and virulence-associated diversity, providing a useful genomic basis for future diagnostic test and vaccine development.

**Data Summary:** Genome assemblies generated for the 153 newly sequenced APP isolates have been deposited at the National Center for Biotechnology Information under BioProject PRJNA1498684. BioSample accessions: SAMN61854423-SAMN61854575, accession numbers and isolate metadata are provided in Supplementary Table S1, and 56 reference sequences used for genotyping are listed in Supplementary Table S2, with the corresponding nucleotide sequences provided in Supplementary Data S2. The overall assembly workflow followed approaches used in previous studies published from our group [1, 2] (https://github.com/isu-vdl-ngs/Brachyspira_hyodysenteriae).

**Impact Statement:** Whole-genome sequencing provides an opportunity to move beyond conventional serotyping and interpret *Actinobacillus pleuropneumoniae* within its broader evolutionary and virulence-associated genomic context. In this study, we integrated population structure, capsular K type, O-antigen type and Apx toxin profiles across a geographically diverse collection of APP genomes. Our findings show that combined K:O antigen characterization provides greater resolution of genomic and toxin diversity than capsular type alone, distinguishing isolates with different genomic backgrounds and toxin profiles that would otherwise be grouped within the same conventional serovar. We also demonstrate that the APP-specific *apxIVA* locus is more structurally diverse than commonly assumed, highlighting the importance of considering population-level sequence variation when interpreting diagnostic and antigenic targets. Together, this work provides a framework for more informative genomic surveillance of APP and supports the future development and evaluation of molecular diagnostics, epidemiological typing schemes and vaccines that account for the broader genetic diversity of circulating APP populations.

## Introduction

*Actinobacillus pleuropneumoniae* (APP) is a gram-negative member of the family *Pasteurellaceae* and the etiological agent of porcine pleuropneumonia, a severe and highly contagious respiratory disease of pigs. Infection can cause rapidly progressive fibrinohemorrhagic and necrotizing pneumonia with substantial morbidity and mortality [3–5]. Disease severity varies among isolates and reflects differences in bacterial virulence determinants together with host and environmental factors [6].

APP pathogenicity is multifactorial and depends on determinants involved in colonization, nutrient acquisition, persistence and tissue damage [4]. These include type IV pili involved in adhesion and biofilm formation [7], transferrin-binding proteins TbpA/TbpB [8], and the haemoglobin receptor HgbA [9]. In contrast to these broadly distributed functions, the Apx repeats-in-toxin (RTX) proteins are major mediators of tissue injury, leukocyte toxicity and hemolysis and are closely associated with differences in virulence among APP isolates [10].

Four Apx toxin systems, ApxI–ApxIV, have been described in APP. ApxI is strongly hemolytic and cytotoxic, ApxII is weakly hemolytic and moderately cytotoxic, and ApxIII is non-hemolytic but strongly cytotoxic [4, 5, 10]. ApxI–ApxIII repertoires vary among isolates, whereas ApxIVA-associated sequence is considered species-associated [11]. Classical toxin typing has established characteristic associations between capsular (K) serotypes and Apx toxin profiles. Serotypes 1, 5, 9, 11, and 16 typically produce both ApxI and ApxII, whereas other serotypes commonly exhibit an ApxII+ApxIII profile. In contrast, serotypes 10 and 14 typically produce ApxI alone, while serotype 13 may produce either ApxI or ApxII alone, depending on geographical origin [12]. Experimental studies have demonstrated major contributions of these toxins to characteristic pulmonary lesions and full virulence [12–14]. These associations have made serotype a practical indicator of expected toxin repertoire, but the relationship is not absolute. Among 213 German clinical isolates, 93.4% showed the Apx profile expected for their serotype and 6.6% showed atypical profiles [11].

ApxIVA is an APP-specific RTX determinant that remains less well characterized than ApxI–ApxIII, in part because of its large coding sequence (>5 kb), repeat-rich architecture and infection-associated expression [14, 15]. Functional studies have shown that ApxIVA contributes to full virulence in experimentally infected pigs [16]. Unlike the classical pore-forming ApxI–ApxIII toxins, ApxIVA contains a Ca²⁺-dependent self-processing module (SPM) that mediates autocatalytic cleavage at a defined Asp–Pro bond followed by covalent cross-linking of the processed N-terminal fragment; deletion of this SPM attenuated APP virulence in a pig infection model [17]. More recently, the N-terminal portion of ApxIVA was shown to possess antibacterial activity, while the small ORF immediately upstream encodes a Cmi-like periplasmic immunity protein, supporting a toxin– immunity function for the bicistronic *apxIVA* operon [18]. Structural variation was already apparent in the original description of ApxIVA, which reported variation in glycine-rich RTX repeat modules among serotypes [19]. Subsequent studies have further demonstrated substantial structural diversity within the *apxIVA* locus, including *ISApl1*-mediated insertions, nonsense mutations, gene fragmentation, and partial *apxIV-S/apxIVS′* duplications associated with mobile genetic elements [15, 20, 21]. Notably, insertion of *ISApl1* within *apxIVA* can disrupt the coding sequence and potentially abolish expression of an intact ApxIVA protein [21]. However, the extent to which complete apxIVA architecture and length variation are structured across the wider APP population remains poorly defined.

APP also exhibits substantial surface-antigen diversity. Capsular polysaccharide (CPS) forms the basis of conventional K-serotype classification [22], and individual K types are often associated with characteristic Apx repertoires [12]. However, the cps locus represents only one component of antigenic diversity. The lipopolysaccharide O-antigen is also variable, and related O-antigen structures can occur across different K types [23]. Isolates sharing the same K type can differ in both O-antigen background and Apx toxin repertoire [24, 25]. K genotyping alone may therefore obscure biologically relevant diversity that can be resolved by combined K and O characterization.

Whole-genome sequencing (WGS) enables antigenic and virulence-associated traits to be interpreted within the evolutionary structure of APP. Comparative analysis of 12 serotype reference genomes identified a broadly conserved genomic backbone with pronounced variability in surface polysaccharide and virulence-associated regions [26]. Subsequent studies showed distinct phylogenetic lineages among serotype 8 isolates from Norway, Denmark and the United Kingdom [27], substantial variation at prophage, CPS, LPS and RTX-toxin regions among 26 complete genomes representing 18 serotypes [28], and geographically structured population diversity among Spanish and publicly available genomes [29]. Genome-based serotyping has also improved resolution of CPS diversity [30, 31]. However, population structure, combined K:O genotype, ApxI–ApxIII repertoire and *apxIVA* structural diversity have not yet been integrated across a comparably broad APP collection.

Here, we analyzed 328 APP genomes, comprising 153 newly sequenced field isolates from United States, Spain and Mexico and 175 publicly available genomes representing 21 countries. We integrated core-genome population structure with K and O antigen genotypes and Apx toxin profiles to test whether K:O genotype provides greater concordance with genomic lineage and ApxI–ApxIII genotype repertoire than K type alone, and to characterize population-level structural diversity of *apxIVA* gene.

## Materials and Methods

### Bacterial isolates and genome collection

A total of 153 APP isolates were obtained from the Iowa State University Veterinary Diagnostic Laboratory (ISU VDL) culture collection. Isolates originated from clinical diagnostic and surveillance submissions received from 2020 to 2025 in the USA (n=124), Spain (n=25) and Mexico (n=4) (Supplementary Table S1). Isolates were recovered from −80 °C frozen stocks and passaged on chocolate agar supplemented with NAD (V factor). Plates were incubated at 38 °C for 48 h in 5% CO₂. Species identification was confirmed by MALDI-TOF MS using the Bruker protocols in routine use at ISU VDL. An additional 175 publicly available APP genome assemblies from 21 countries were included for comparative analysis (Supplementary Table S1), resulting in a final dataset of 328 genomes.

### DNA extraction, sequencing and assembly

Genomic DNA was extracted from bacterial pellets using the MagMAX Pathogen RNA/DNA Kit on a KingFisher Flex System (Thermo Fisher Scientific, Waltham, MA, USA) according to the manufacturer’s instructions. DNA concentration was measured using a Qubit fluorometer with the Qubit 1× dsDNA HS Assay Kit (Life Technologies, Carlsbad, CA, USA). Indexed libraries were prepared using the Nextera XT DNA Library Preparation Kit (Illumina, CA, USA), pooled, and sequenced on an Illumina MiSeq platform using MiSeq Reagent Kit v2 (500 cycles) to generate approximately 2 × 250-bp paired-end reads.

Raw-read quality was assessed using FastQC and summarized with MultiQC [32]. Low-quality sequence and adapters were trimmed with Trimmomatic v0.36 [33], followed by repeat FastQC/MultiQC assessment. De novo assemblies were generated with SPAdes v3.15.5 [34] using the --careful option. Assembly quality was assessed with QUAST [35]. Across the dataset, genome sizes were consistent with APP and GC content was approximately 41%.

### Core-genome phylogeny and population structure

All 328 assemblies were annotated with Prokka v1.14.6 [36] and analysed with Panaroo v1.3.4 to construct the pangenome and core-gene alignment. Variable sites from core gene clusters present in ≥99% of genomes were extracted with SNP-sites v2.5.1 [37]. A maximum-likelihood tree was reconstructed from the variable-site alignment with IQ-TREE v2.1.4, and ModelFinder selected the SYM+ASC+R4 by Bayesian information criterion [38, 39]. The +ASC correction was applied because the phylogenetic alignment contained variable sites only, and branch support was assessed with 1,000 ultrafast bootstrap replicates. FastBAPS v1.0.8 [40] was applied to the core-genome SNP alignment using the baps option.

### Capsule K and O-antigen genotyping

Capsular K types were assigned by BLASTn comparison against a curated reference database containing representative cps loci for the 19 recognized capsular types (Supplementary Table S2, Supplementary Data S2). The final K type was assigned from the best complete-locus match meeting ≥95% nucleotide identity and ≥95% reference coverage. Assignments were independently cross-checked with Serotype Detector [31], which evaluates cps gene composition and sequence similarity against reference-serotype profiles.

O-antigen types were assigned using a custom nucleotide database representing the O-antigen-specific loci targeted by the multiplex-PCR scheme of To et al. [24] (Supplementary Table S2, Supplementary Data S2). Depending on O type, targets included loci within *wcaJ*, *wbaP* and glycosyltransferase/acyltransferase genes. An O type was assigned when the best reference match met ≥95% nucleotide identity and ≥95% coverage. K and O assignments were then combined and reported in K:O format.

### Detection of virulence-associated genes and ApxI–ApxIII genotyping

Virulence-associated loci were screened with ABRicate against VFDB and a species-focused custom database assembled from published APP sequences (Supplementary Table S2, Supplementary Data S2). The custom database included genes involved in iron acquisition, adhesion and colonization, biofilm formation, outer membrane structure and RTX toxin systems, including *hgbA*, *tbpA*, *tbpB*, *afuABC*, *fhuB*, *pilW*, *pilQ*, *pil*, *apf*, *ashA*, *ashC*, *pal*, *ompA*, *pgaABCD*, *flp* and the Apx toxin loci. Homologues were considered detected at ≥90% nucleotide identity and ≥90% sequence coverage. Activation gene (*apxIC*, *apxIIC*, *apxIIIC*), structural toxin gene (*apxIA*, *apxIIA*, *apxIIIA*), and corresponding secretion-associated genes (*apxIB*, *apxIIIB*, *apxID*, *apxIIID*) were evaluated separately for the Apx genotypes. An isolate was classified as ApxI genotype positive when the *apxICABD* was present; ApxII genotype positive when both *apxIICA* and *apxIBD* were present; and ApxIII genotype positive when the *apxIIICABD* operon was present [41–43]. Because threshold-based screening can miss highly divergent alleles or fragmented loci, non-detection was not interpreted as definitive gene absence without locus-level confirmation.

### Comparative analysis of apxIVA

The apxIVA locus was analyzed separately because of its large size, repeat-rich RTX architecture and frequent fragmentation in draft assemblies. All 328 genomes were screened by BLASTn against five representative apxIVA sequences from serotypes 1, 2, 3, 5 and 8 (Supplementary Table S2, Supplementary Data S2), using a minimum nucleotide identity of 90% and a minimum aligned length of 200 bp for initial detection. Complete single-contig loci were defined as hits with ≥95% identity and ≥95% reference coverage. Initial screening identified 62 genomes with a complete single-contig apxIVA open reading frame. Six of these genomes lacked the required 10-kb upstream and downstream flanking regions and were excluded; consequently, structural comparison was restricted to 56 genomes containing an uninterrupted complete apxIVA open reading frame on a single contig together with sufficient flanking sequence. For each retained genome, the apxIVA coding sequence and 10 kb of upstream and downstream sequence were extracted. Gene organization and local synteny were compared with Clinker, and predicted ApxIVA proteins were aligned with MAFFT and inspected in UGENE. Length variation was described relative to the ApxIVA protein encoded by GCF_021957185.1, with alignment gaps reported as reference-relative differences rather than inferred evolutionary deletions. The 56 complete loci constituted a quality-selected subset and were not used to estimate lineage-specific prevalence of structural classes.

### Statistical analysis

Directional concordance among capsular K type, combined K:O genotype, FastBAPS lineage and Apx genotype was quantified using the adjusted Wallace coefficient (AW) [44, 45]. Comparisons were performed for K→lineage, K:O→lineage, K→Apx genotype, K:O→Apx genotype and lineage→Apx genotype. Ninety-five per cent confidence intervals were estimated from 10,000 non-parametric bootstrap resamples of isolates. For pairwise comparison of two AW coefficients, the same resampled isolate indices were used for both coefficients and the paired difference was calculated in each bootstrap replicate. A difference was considered supported when the 95% bootstrap confidence interval of the paired difference excluded zero. A fixed random seed of 20260727 was used for reproducibility. These statistics quantify concordance within the sampled collection and do not establish causal or independent evolutionary effects.

## Results

### Whole-genome population structure and geographical distribution

The final dataset comprised 328 APP genomes: 153 newly sequenced isolates from the United States (n=124), Spain (n=25) and Mexico (n=4), and 175 publicly available genomes from around the world. Newly sequenced assemblies were 2.2–2.3 Mbp in length, had a median sequencing depth of 55× and a GC content of approximately 41%. The collection represented 21 countries across four continents, although sampling was markedly uneven. The largest country-level contributions were from the USA (n=139), Spain (n=42), Canada (n=25), China (n=23), Switzerland (n=21), Denmark (n=15) and Germany (n=14) (Supplementary Table S1).

Panaroo identified 4,335 pangenome gene clusters, including 1,818 core genes present in ≥99% of genomes. The core-gene alignment contained 146,902 variable SNP sites. IQ-TREE reconstructed the maximum-likelihood phylogeny under the SYM+ASC+R4 model selected by ModelFinder, with 1,000 ultrafast bootstrap replicates [38, 39]. FastBAPS partitioned the core-genome SNP alignment into 23 clusters, hereafter referred to as genomic lineages L1–L23 (Fig. 1).

**Fig. 1.**
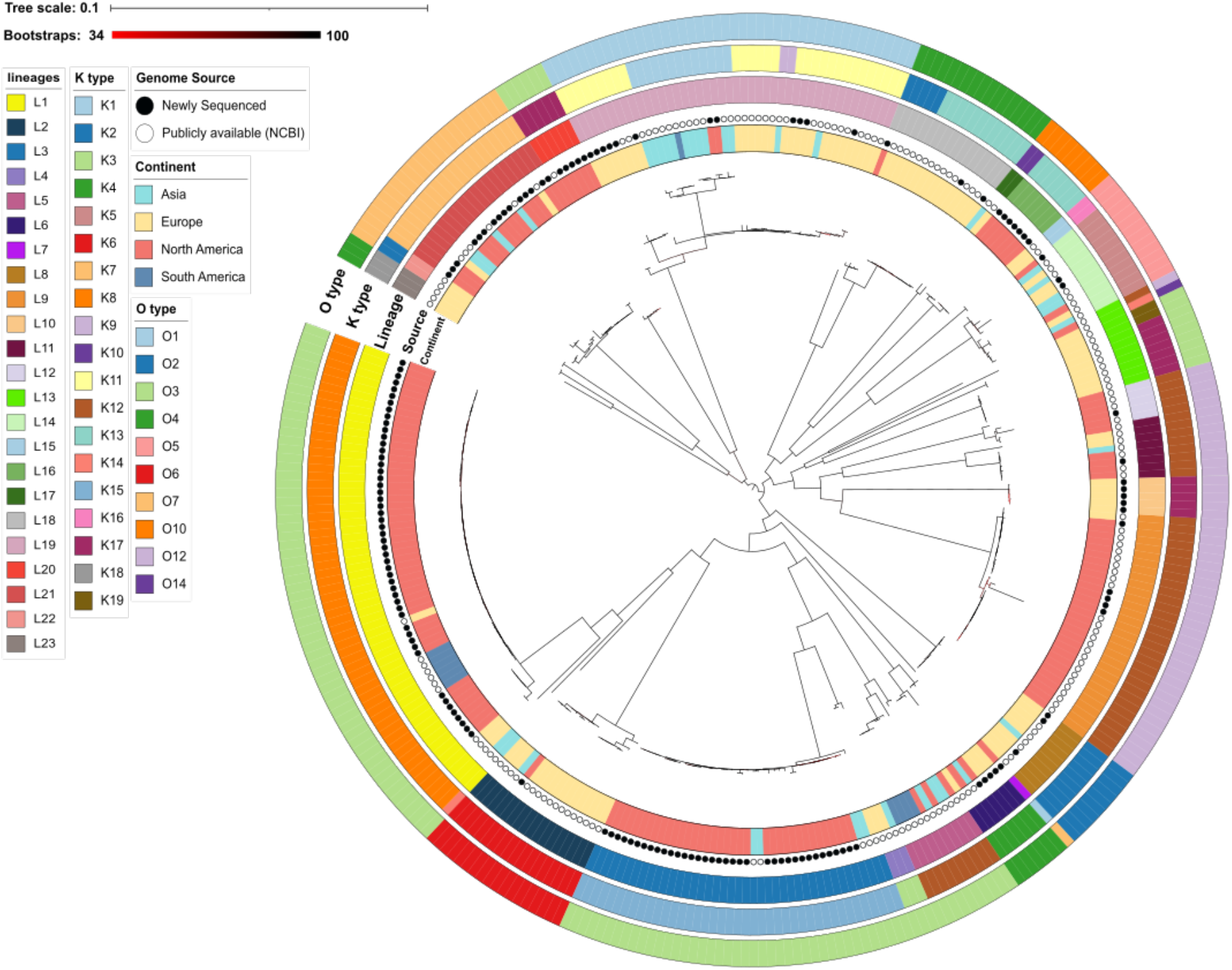
Core-genome phylogeny and genomic characteristics of 328 Actinobacillus pleuropneumoniae genomes. A midpoint-rooted maximum-likelihood phylogeny was reconstructed from 146,902 variable core-genome SNP sites using IQ-TREE v2.1.4 under the SYM+ASC+R4 model. Annotations indicate continent, genome source, FastBAPS genomic lineage (L1–L23), capsular K genotype and O-antigen genotype. Filled and open circles denote genomes newly sequenced in this study and publicly available genomes, respectively. Branch colour intensity represents ultrafast bootstrap support (34–100%), and branch lengths represent nucleotide substitutions per site.

L1 was the largest lineage (63/328, 19.2%), followed by L19 (44/328, 13.4%) and L3 (41/328, 12.5%). Among L1 genomes, 48/63 (76.2%) originated from the USA; 31/44 L19 genomes (70.5%) were European; and 36/41 L3 genomes (87.8%) originated from the USA. These distributions describe the composition of the sampled collection and should not be interpreted as estimates of regional lineage prevalence.

Lineage composition differed among sampled countries (Fig. 2). US genomes were distributed primarily among L1 (48/139, 34.5%), L3 (36/139, 25.9%), L21 (15/139, 10.8%) and L9 (14/139, 10.1%). Twenty-three L3 genomes originated from a single Iowa outbreak during 2021–2022, resulting in substantial epidemiological clustering within this lineage. Chinese genomes occurred in L1–L4, L6, L8, L11, L14, L17, L19 and L21, while Swiss genomes spanned 15 of 23 lineages. In contrast, Canadian, German and Spanish genomes were concentrated in L9 (16/25, 64.0%), L2 (13/14, 92.9%) and L19 (23/42, 54.8%), respectively.

**Fig. 2.**
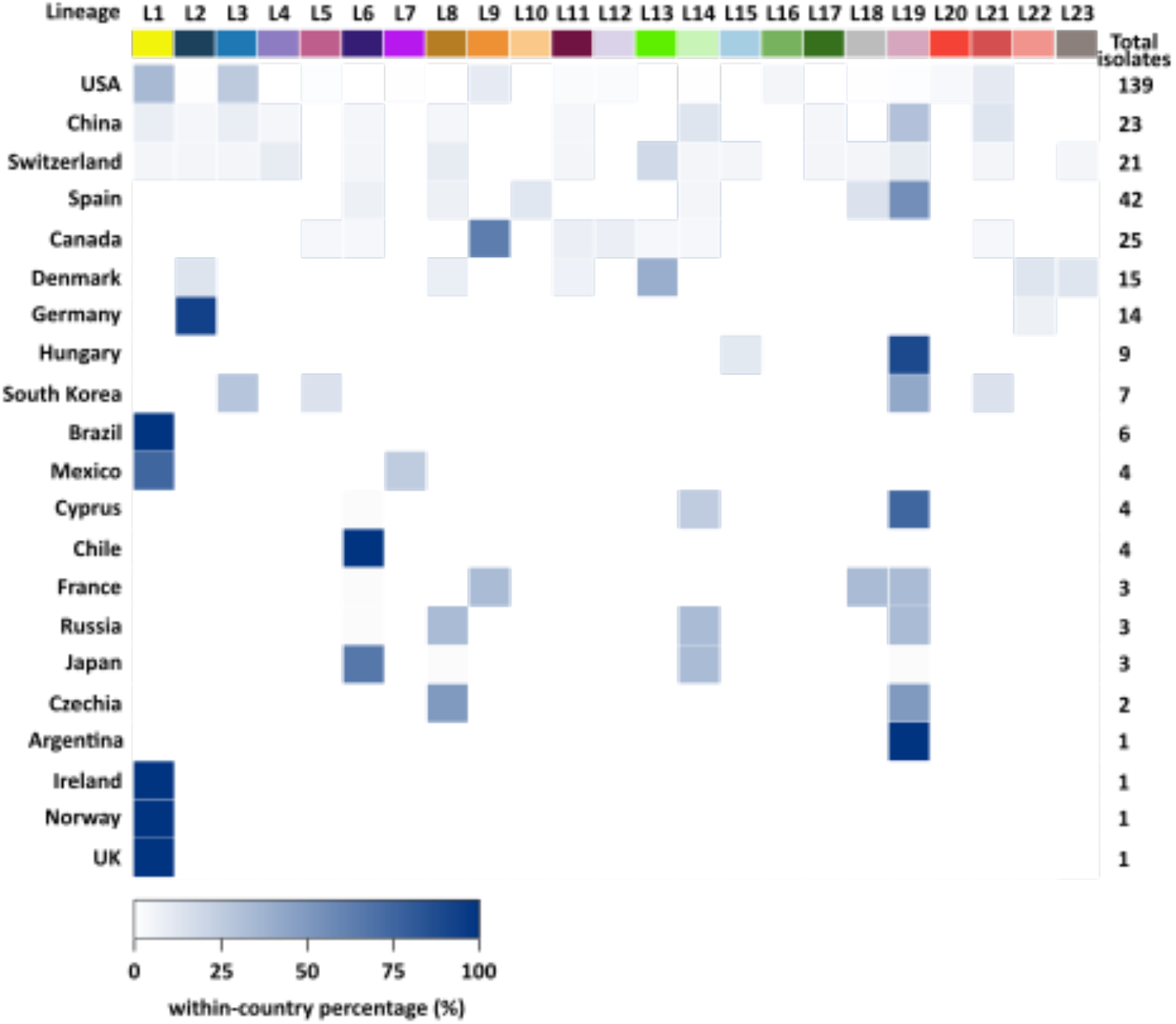
Geographic distribution of genomic lineages in the 328-genome collection. The heatmap shows the percentage contribution of each lineage within each sampled country, with darker shading indicating a higher within-country percentage. The rightmost column gives the total number of genomes sampled from each country. Percentages are descriptive of this dataset; values for countries represented by small numbers of genomes should not be interpreted as population prevalence estimates.

### K:O antigen genotype shows greater concordance with genomic lineage than K type alone

Across the 328 genomes, 19 K types and 10 O-antigen types were identified. K types were assigned by BLASTn screening against the curated cps database (Supplementary Tables S2) and evaluated independently with serotype Detector (Supplementary Table S3) [31]. Serotype Detector output was compatible with the final K-type assignment for all 328 genomes, comprising 231 exact, 60 tentative-compatible and 37 ambiguous-compatible results.

K8 was the most frequent K type (62/328, 18.9%), followed by K12 (55/328, 16.8%), K15 (41/328, 12.5%), K11 (29/328, 8.8%) and K7 (21/328, 6.4%). Six K types occurred in more than one lineage: K1, K2, K12, K13, K14 and K17. K12 showed the broadest lineage distribution and occurred in L5, L9, L11, L12 and L13.

O-antigen types were assigned using reference sequences corresponding to the multiplex-PCR targets described by To et al. [24]. O3 was the most frequent O type (132/328, 40.2%), followed by O12 (50/328, 15.2%), O1 (44/328, 13.4%), O4 (27/328, 8.2%) and O7 (24/328, 7.3%). Only O1, O2, O6 and O14 were each confined to a single lineage in this dataset.

Combining K and O assignments identified 26 K:O genotypes, of which 24 (92.3%) were restricted to a single lineage (Table 1). The exceptions were K12:O12, which occurred in L9 and L11–L13, and K17:O3, which occurred in L13 and L20. K8:O3, restricted to L1, was the most frequent genotype (62/328, 18.9%), followed by K12:O12 (45/328, 13.7%) and K15:O3 (41/328, 12.5%), the latter restricted to L3.

**Table 1.**
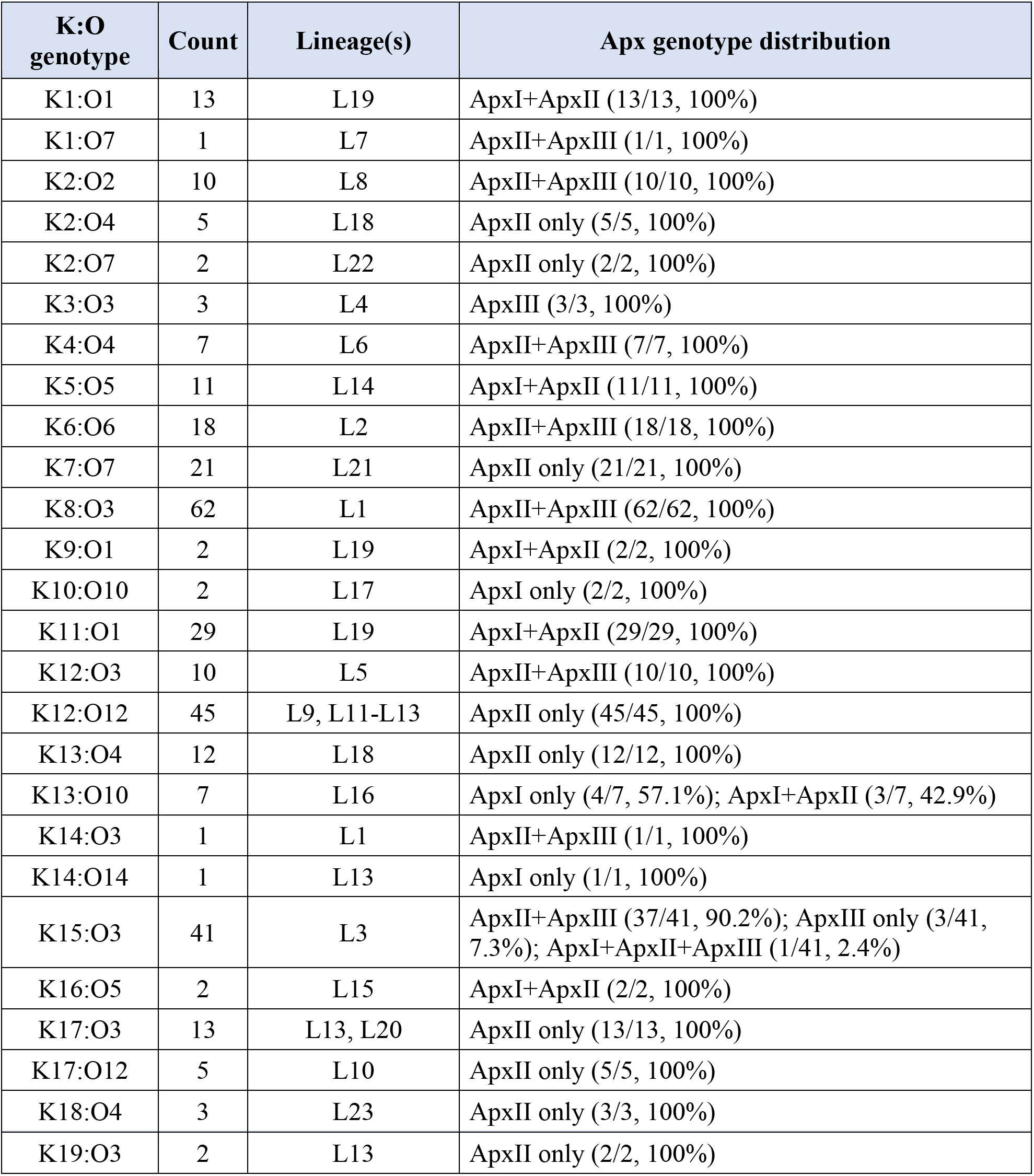
K:O antigen genotypes, genomic lineages and ApxI–ApxIII genotype distributions. An isolate was classified as ApxI genotype positive when the *apxICABD* was present; ApxII genotype positive when both *apxIICA* and *apxIBD* were present; and ApxIII genotype positive when the *apxIIICABD* operon was present.

O-antigen assignment subdivided several K types into phylogenetically distinct groups. K1:O1 was restricted to L19 whereas K1:O7 occurred in L7. K2:O2, K2:O4 and K2:O7 occurred in L8, L18 and L22, respectively. K13:O4 was restricted to L18 whereas K13:O10 was restricted to L16. Similarly, K14:O3 and K14:O14 occurred in L1 and L13, respectively, while K17:O12 occurred in L10 and K17:O3 in L13 and L20.

Directional concordance analysis supported closer correspondence between K:O genotype and population structure. The adjusted Wallace coefficient for K:O→lineage was 0.881 (95% bootstrap CI, 0.814–0.948), compared with 0.759 for K→lineage (95% CI, 0.672–0.847). The paired difference was 0.122 (95% CI, 0.072–0.172), supporting greater concordance with genomic lineage after incorporation of O-antigen type [44]. These associations are specific to the sampled collection and do not demonstrate an independent evolutionary effect of O-antigen type on lineage structure.

### Non-Apx virulence-associated loci

Initial screening against VFDB recovered only *gmhA*. A species-focused database was therefore used to screen 29 loci associated with iron acquisition, adhesion, biofilm formation, outer-membrane structure and RTX toxins. Homologues were called at ≥90% nucleotide identity and ≥90% sequence coverage (Supplementary Table S2).

Most non-Apx loci were detected in nearly all genomes (Fig. 3). The iron-acquisition loci *hgbA* and *fhuB* were detected in 100% of genomes, while *tbpA* and *afuABC* were detected in 99.7%. In contrast, *tbpB* met the detection threshold in 31.7% of genomes and showed strong lineage structure: it was detected in all genomes assigned to L2 and L9–L15, in 29.5% L19 genomes, and not detected at the specified threshold in the remaining lineages. Adhesion-associated loci *pilW, pilQ, pil, apf and ashC* were detected in 97.0–100% of genomes. Gene *pal* and *ompA* were detected in 100% and 99.4%, respectively, while *pgaABCD* and *flp* were detected in 93.9% and 90.2%. Because the screen was threshold-based, the *tbpB* pattern cannot by itself distinguish true absence from divergent or fragmented alleles.

**Fig. 3.**
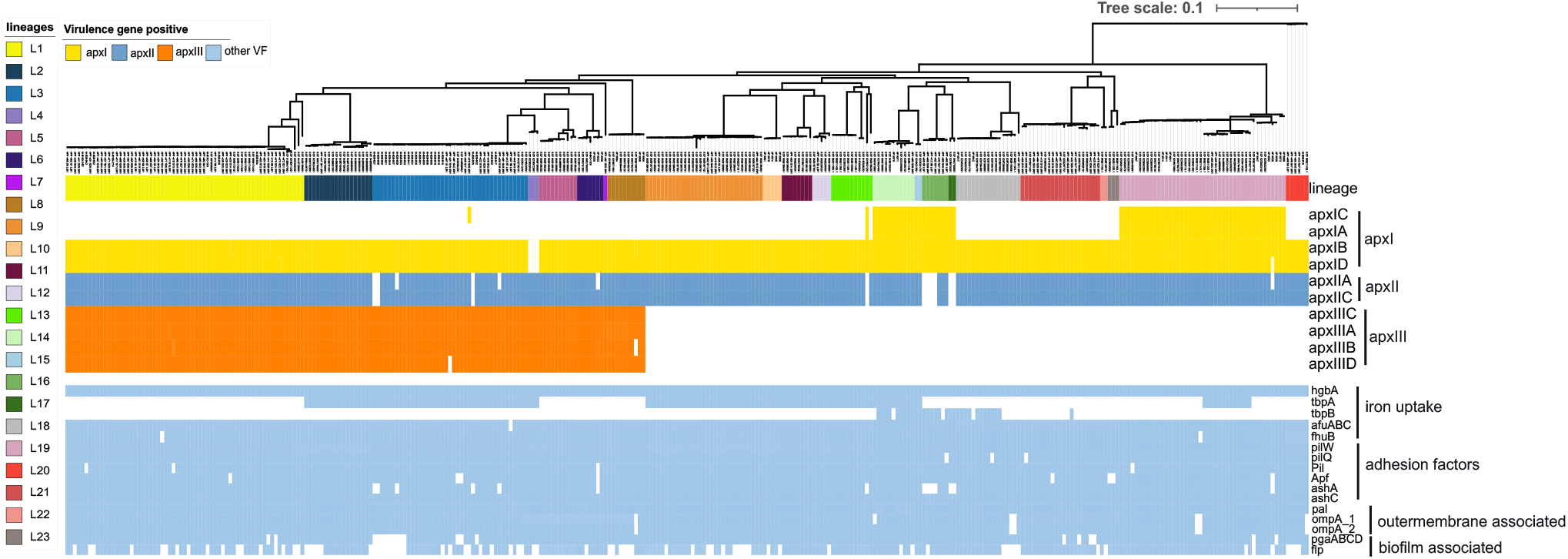
Virulence-associated gene profiles across genomic lineages. The upper annotation bar indicates the 23 FastBAPS lineages. The matrix shows loci meeting the detection threshold (≥90% nucleotide identity and ≥90% sequence coverage) for ApxI–ApxIII components, iron-acquisition systems, adhesion factors, outer-membrane-associated loci and biofilm-associated loci.

### ApxI–ApxIII genotypes co-segregate with lineage and K:O background

ApxI, ApxII, and ApxIII genotypes were defined as positive only when the complete corresponding toxin gene sets were present: *apxICABD* for ApxI, *apxIICA* together with *apxIBD* for ApxII, and *apxIIICABD* for ApxIII. Six genotype combinations were observed: ApxII+ApxIII (146/328, 44.5%), ApxII only (108/328, 32.9%), ApxI+ApxII (60/328, 18.3%), ApxI only (7/328, 2.1%), ApxIII only (6/328, 1.8%) and ApxI+ApxII+ApxIII (1/328, 0.3%) (Table 1).

ApxIII-positive genotypes occurred in 153/328 genomes (46.6%) and were confined to L1–L8. ApxI-positive genotypes occurred in 68/328 genomes (20.7%); 66/68 were in L14–L17 or L19, with one additional genome in L3 and one in L13 (Fig. 3). ApxII-positive genotypes occurred in 315/328 genomes (96.0%). The shared ApxI/ApxII secretion genes *apxIB* and *apxID* were detected in 325/328 genomes but were not detected in three L4 K3:O3 genomes. These three genomes nevertheless contained paired ApxII activation and structural genes, however, genomic detection of *apxIICA* alone does not establish functional toxin secretion.

Apx genotype was homogeneous within 24 of the 26 K:O groups (Table 1). Several K types contained distinct Apx genotypes after subdivision by O type. K1:O1 carried ApxI+ApxII whereas K1:O7 carried ApxII+ApxIII; K2:O2 carried ApxII+ApxIII whereas K2:O4 and K2:O7 carried ApxII only; and K14:O14 carried ApxI only which is consistent with traditionally reported K14 whereas K14:O3 carried ApxII+ApxIII. The exceptions were K13:O10, which comprised four ApxI-only and three ApxI+ApxII genomes, and K15:O3, which comprised 37 ApxII+ApxIII which was traditionally described in K15, three ApxIII-only and one ApxI+ApxII+ApxIII genome.

K:O genotype showed high directional concordance with Apx genotype (adjusted Wallace coefficient, 0.949; 95% bootstrap CI, 0.899–0.989), compared with 0.787 for K type alone (95% CI, 0.705–0.871). The paired difference was 0.163 (95% CI, 0.090–0.239). Genomic lineage showed similarly high concordance with Apx genotype (0.947; 95% CI, 0.897–0.987), and did not differ detectably from K:O genotype (paired difference, 0.002; 95% CI, −0.005 to 0.011) [44]. Thus, Apx genotypes co-segregated closely with both genomic lineage and lineage-associated K:O genotype. Because 24/26 K:O genotypes were restricted to individual lineages, these analyses do not separate independent effects of lineage and antigen genotype on Apx composition.

Geographic differences in *apxCA* profiles largely mirrored the unequal geographic distribution of lineages and K:O genotypes. Within the sampled collection, *apxICA*-containing profiles contributed a larger share of detected *apxCA* loci in several European and Asian country subsets, whereas US genomes were dominated by *apxIICA* and *apxIIICA*-containing profiles (Fig. 4). Because country sample sizes were highly unequal and several countries were represented by very few genomes, these patterns are descriptive and should not be interpreted as regional prevalence estimates.

**Fig. 4.**
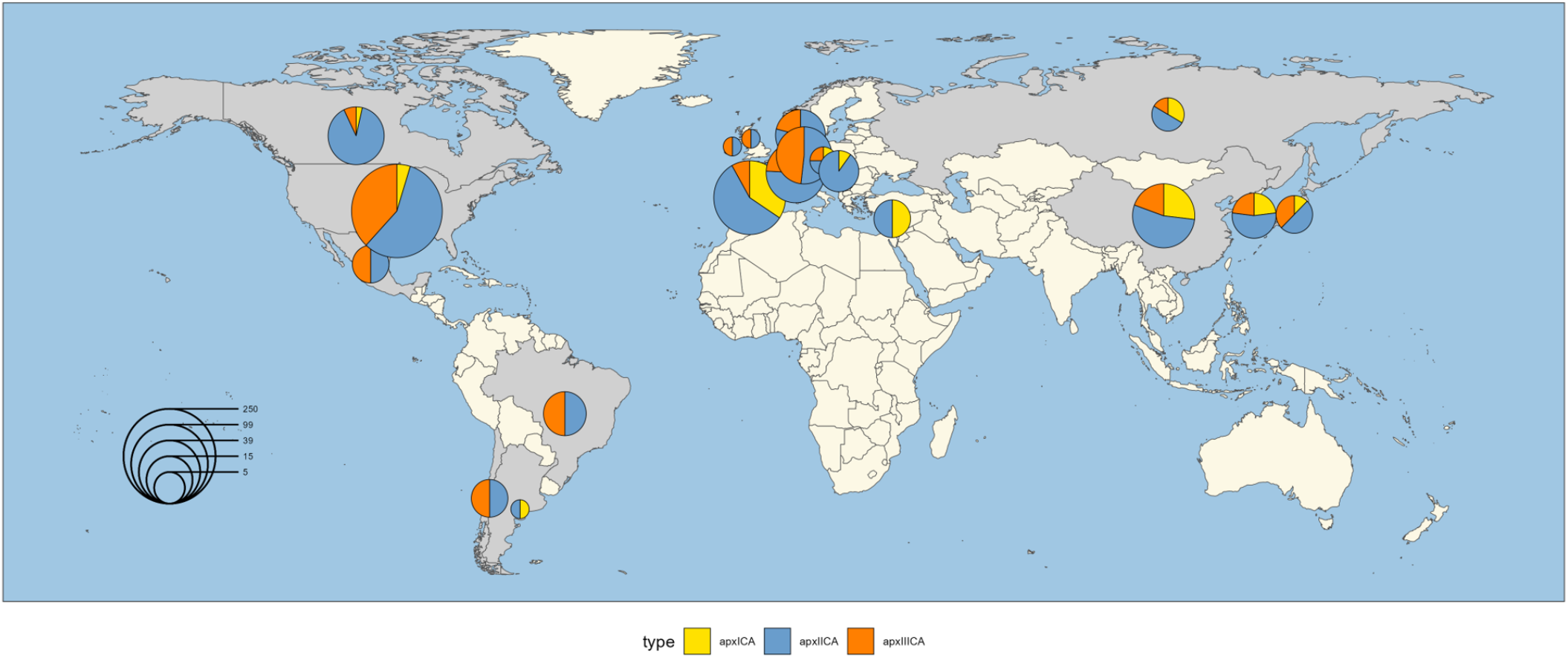
Geographic distribution of *apxCA* locus counts in the sampled collection. Pie sectors represent detected *apxICA, apxIICA* and *apxIIICA* loci among genomes from each country. Circle area is scaled to the total number of detected *apxCA* loci and therefore reflects sampling intensity as well as genotype composition. The map is descriptive and should not be interpreted as an estimate of regional Apx prevalence.

### Lineage-associated *apxIVA* length variation and *apxIVA*-related homologs

BLASTn screening detected apxIVA-related sequence across the 328-genome collection. Multiple non-overlapping hits were common in draft assemblies, consistent with fragmentation of this long, repeat-rich locus across contigs. Initial screening identified 62 complete single-contig apxIVA open reading frames; six lacked sufficient 10-kb upstream and downstream flanking regions and were excluded from the flanking-region analysis. The final structural comparison therefore included 56 complete single-contig loci representing 15 lineages. Coding lengths ranged from 4,149 to 5,841 bp. A 474-bp ORF reported to encode a Cmi-like periplasmic immunity protein protecting cells from the antibacterial activity of ApxIVA [18] was located immediately upstream of apxIVA in all 56 loci. In some genomes, an additional ApxIVA-related homolog of variable length occurred between the apxIVA locus and lacZ together with an IS256-family transposase and a hypothetical protein. Core gene order within the 10-kb flanking regions was otherwise conserved (Fig. 5). Because complete single-contig loci were unevenly represented among lineages, structural-class counts are descriptive rather than prevalence estimates.

**Fig. 5.**
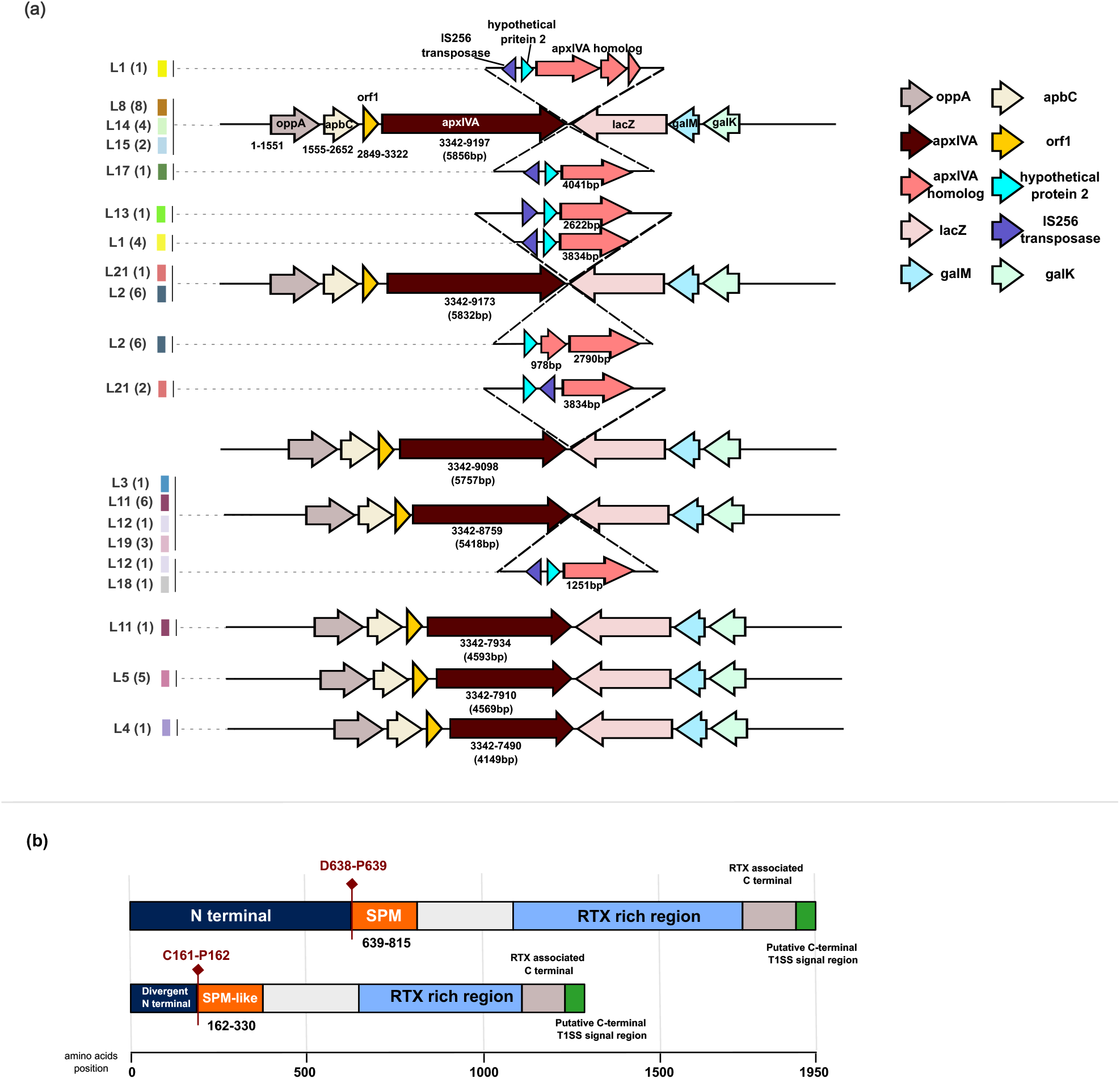
Structural variation of the apxIVA locus and ApxIVA-related homolog. (a) Representative apxIVA locus architectures and flanking gene organization among the 56 complete single-contig apxIVA loci from 15 genomic lineages. Architectures are grouped by recurrent reference-relative length pattern. Numbers in parentheses indicate the number of genomes represented within the corresponding lineage/architecture group; triangles denote genomes containing an additional ApxIVA-related homolog. (b) Schematic comparison of predicted functional regions in canonical ApxIVA and the divergent ApxIVA-related homolog, including the experimentally defined SPM, RTX repeat-rich region, C-terminal secretion-associated region and the divergent SPM-like segment in the homolog.

A 1,277-aa protein represented by GCF_019264705.1 was consistent with an ApxIVA-related homolog (Fig. 5). Sequence conservation was highest in the putative C-terminal RTX-associated region (∼92%). The predicted RTX repeat-rich region shared approximately 82% amino-acid identity with the corresponding canonical ApxIVA region, and an RTX calcium-binding repeat region corresponding to PF00353 showed approximately 90% identity. By contrast, the putative SPM-like region of the homolog (aa 162–330) shared only 37.9% identity with the experimentally defined ApxIVA SPM (aa 639–815). The homolog also lacked the canonical D638–P639 autocleavage boundary, with C161– P162 at the corresponding aligned position. Calcium-dependent clip-and-link activity therefore cannot be inferred for this homolog from sequence similarity alone.

Alignment of the 56 predicted ApxIVA proteins to GCF_021957185.1 identified recurrent reference-relative length patterns (Supplementary Table S4). Sixteen proteins (28.6%) contained no alignment gaps relative to the reference, 20 (35.7%) contained only N-terminal reference-relative differences, 13 (23.2%) contained a 146-aa reference-relative gap within the RTX repeat-rich region, and seven (12.5%) contained a 421-aa gap in the same region. All 12 L2 proteins shared an 8-aa N-terminal reference-relative gap, while all five L5 proteins combined this 8-aa difference with the 421-aa RTX-region gap. Breakpoints were highly conserved among the larger variants: 12/13 proteins in the 146-aa group lacked residues corresponding to reference positions 1151–1296, and 6/7 in the 421-aa group lacked positions 1151–1571. The experimentally defined SPM and canonical D638–P639 processing boundary were retained in all 56 canonical ApxIVA proteins.

## Discussion

WGS has increasingly been used to characterize APP, but previous genomic studies have generally focused on reference strains, individual serotypes, geographically restricted collections or selected aspects of antimicrobial resistance and virulence [4, 26–29]. This study integrates 153 newly sequenced isolates with 175 publicly available genomes representing from 21 countries and places, K and O antigen genotypes, ApxI–ApxIII repertoire and ApxIVA architecture within a common population-genomic framework.

Overall, the toxin structural and secretion genes showed highly consistent co-occurrence across the dataset. *apxICA* was consistently accompanied by *apxIBD*, *apxIIICA* by *apxIIIBD*, and *apxIICA* by *apxIBD*, supporting the conserved organization of these toxin–secretion gene modules. A notable exception was observed in lineage L4, in which all three isolates carried *apxIICA* but lacked *apxIBD*. Even though all three *apxIB* and *apxID* sequences showed similarly high coverage of 99.48% and 98.33%, respectively, both loci exhibited consistently low sequence identity, at 75.53% for *apxIB* and 71.69% for *apxID*. The consistent absence of *apxIBD* in isolates from this lineage, despite retention of the ApxII structural and activator genes, suggests that ApxII secretion may be changed or impaired or may rely on an alternative secretion system. These possibilities require further experimental validation.

Previous classical molecular studies established an association between APP K serotype and Apx toxin repertoire [12, 46], and field collections have generally retained these relationships. For example, 199/213 German clinical isolates had the Apx profile expected for their serotype [11], while a recent Korean survey found the classical ApxI+ApxII configuration in all 88 serotype 1 isolates examined [47]. Our data were broadly consistent with these established associations. For example, among 14 K1 isolates, 13 carried the classical ApxI+ApxII profile, whereas one remaining isolate, classified as K1:O7, carried an atypical ApxII+ApxIII profile and differed from the other K1 isolates in both O-antigen genotype and phylogenetic lineages. This exception illustrates a limitation of conventional locus-based typing: although capsule and toxin markers can capture characteristic serotype-associated patterns, they do not resolve the broader genomic background in which these antigen and toxin loci occur. Whole genome analysis therefore provides additional context for characterizing individual antigen and toxin loci among genome-wide related isolates.

More recent WGS studies have demonstrated associations among serotype, Apx profile, and phylogenetic clustering, highlighting the additional resolution provided by genomic context. Some reference-genome representing 18 serotypes comparative study have been analyzed, showing genome-wide phylogeny and surface-antigen diversity [26, 28, 48], but essentially not population genomics instead of one or two reference strains per serotype. Cohen conducted comparative genomic analysis of 212 serotype 8 isolates from Norway, Denmark and the UK and revealed the same serotype can contain deeply separated genomic lineages [27]. Similar findings were also shown in 169 Spanish clinical isolates that the same serotypes contained more than one clonal group [49]. A most recent study of 12 Italian isolates also identified distinct genomic lineages within serotype 6 [50]. However, those studies have largely focused on reference genomes, individual serotypes or geographically restricted populations. Our present study included a large population from 21 countries and all currently circulating 19 serotypes, and further revealed that the same serotype from different geographical regions can be clustered into different clonal groups.

The relationship between capsule and O-antigen variations helps explain why K type alone does not fully capture APP genomic diversity [4]. Structural studies showed that O antigens can be shared across different capsular types [23]. More recent work identified K2:O4 and K2:O7 variants and established genomic targets for O-antigen typing [24], and K6b:O3 has subsequently been described [51]. In the present collection, 26 K:O genotypes were identified and 24 were restricted to a single genomic lineage. O typing separated several K types into distinct genomic groups, including K1:O1 versus K1:O7, K2:O2/K2:O4/K2:O7, K13:O4 versus K13:O10 and K14:O3 versus K14:O14. Quantitatively, K:O showed greater directional concordance with lineage than K alone, supporting combined antigen genotyping as a higher-resolution descriptor of the sampled population structure.

Historical classifications linked K types 1, 5, 9, 10 and 11 with higher virulence and K types such as 3, 7 and 12 with lower virulence, largely on the basis of characteristic Apx profiles [52]. Atypical field isolates have nevertheless been documented; Schuwerk et al. detected non-classical Apx profiles in 6.6% of German isolates, including K types 2 and 7 exceptions [11]. Particularly relevant one is the study by Vincent et al., in which atypical K type 12 strains carried ApxII+ApxIII rather than ApxII alone, had a distinct O-antigen background and phylogenetic placement, and caused more severe disease than the classical K type 12 reference strain in pigs [25]. These observations support evaluating toxin repertoire within broader genomic and antigenic context rather than inferring pathogenic potential from capsule type alone. Our data further resolved the relationships among K:O genotypes, Apx genotypes and phylogenetic groups. Phylogenetic lineages provide greater concordance with ApxI-ApxIII repertoire than K type alone. K:O genotypes showed similarly high concordance with Apx genotype, and did not differ detectably from lineages. For example, all K2:O2 from L8 genomes carried ApxII+ApxIII, while K2:O4 from L18 and K2:O7 from L22 carried ApxII only. K type-associated Apx genotype patterns remain useful for anticipating toxin repertoire at the population level, while exceptions are biologically informative and require explanations within genomic context. This finding has important practical implications for diagnostics and surveillance, and potentially for vaccine development. With multiplex PCR methods for comprehensive O-antigen typing now available [24], combining K and O typing could provide a practical and relatively inexpensive approach for improving strain discrimination, particularly in diagnostic laboratories where routine WGS of large numbers of isolates may not be feasible. However, we should emphasize that K and O typing is complementary to, rather than a replacement for, WGS. K and O typing examines only two antigenic loci and therefore captures only a small portion of genomic diversity among isolates.

This study included 23 outbreak-associated isolates recovered from Iowa between October 2021 and February 2022, all of which belonged to lineage L3 and carried the K15:O3 genotype. Most outbreak-associated isolates exhibited the canonical ApxII+ApxIII genotype, whereas a small subset (n=3) carried ApxIII alone. Notably, five additional L3/K15:O3 isolates with the ApxII+ApxIII genotype were identified in Switzerland (1), South Korea (2), and China (2), indicating that this was probably not restricted to the Iowa outbreak. It was subsequently detected in U.S. diagnostic submissions during 2024–2025, including isolates from Iowa (n=7), Ohio (n=4), and Indiana (n=2), suggesting continued circulation of this L3/K15:O3 in the United States.

In contrast, the predominant lineage among U.S. isolates sampled during 2024–2025 was L1 (n=44), characterized by the K8:O3 genotype and a uniformly conserved ApxII+ApxIII genotype, indicating that much of the sampled US population was represented by two distinct O3-associated genomic backgrounds with similar toxin repertoires. L1 was also identified in geographically diverse populations, including isolates from Brazil, with earliest isolate collected in 2006, and from Mexico and China. Furthermore, the two oldest genomes in the present NCBI genome collection, recovered from Ireland in 1984 and the United Kingdom in 1995, were assigned to L1. Together, these observations indicate that L1/K8:O3 represents a longstanding and geographically widespread genomic lineage that has persisted across multiple decades and regions.

At the broad population level, the geographic structure of the APP population was characterized by both regional expansion of predominant lineages and the distribution of individual lineages across national boundaries. In the USA, the sampled population was dominated by L1/K8:O3 (48/139, 34.5%) and L3/K15:O3 (36/139, 25.9%) with canonical ApxII and ApxIII genotypes. Additional US populations included L9/K12:O12 and L21/K7:O7, both associated with ApxII alone. Notably, L9 provided a clear example of a lineage shared across neighboring production systems: 30/31 L9 genomes primarily originated from the USA (14/31) and Canada (16/31). Although the present genomic dataset cannot reconstruct transmission routes, this geographic overlap is consistent with dissemination through interconnected swine populations and transboundary animal movement between the two countries [53], illustrating how regional production networks may contribute to the maintenance of shared APP clones. In contrast, European populations showed greater geographic differentiation in dominant genomic backgrounds. L19 was strongly represented in Europe, including 23/42 Spanish genomes, and was associated with O1-containing K:O genotypes (44/44) and ApxI+ApxII configuration (42/44), whereas Germany was dominated by L2/K6:O6 (13/14, 92.9%), which carried ApxII+ApxIII, and Denmark contained a substantial L13 population (6/15, 40.0%), predominantly associated with ApxII-only genotypes. Thus, the geographic distribution of Apx profiles appears to arise largely from the unequal distribution of lineage–K:O. This has potential implications for clinical surveillance: the predominance of L1/K8:O3 and L3/K15:O3 in the US collection suggests that ApxII+ApxIII-associated populations warrant continued monitoring in North America, whereas the substantial representation of L19 and other ApxI-containing backgrounds in Europe supports surveillance for persistence, expansion, or introduction of ApxI-associated lineages. More broadly, monitoring changes in lineage–K:O–Apx combinations may provide greater epidemiological resolution than capsular serotyping alone by helping identify clonal expansion, transboundary dissemination, or replacement of established populations. However, these geographic patterns reflect the composition of the available genome collection and should not be interpreted as estimates of national prevalence, direct evidence of transmission, or evidence that any particular Apx genotype is intrinsically associated with greater disease severity.

Most non-Apx virulence-associated loci were broadly conserved, consistent with functions maintained across diverse APP genomic backgrounds. *hgbA, fhuB, tbpA* and *afuABC* were detected in nearly all genomes, as were most adhesion and biofilm-associated loci. *tbpB* was an exception, meeting the screening threshold in only 31.7% of genomes and showing strong lineage structure. Because this conclusion derives from nucleotide identity and coverage thresholds, apparent *tbpB* non-detection may reflect true absence, substantial allelic divergence or local assembly fragmentation. Locus-level analysis would therefore be required before interpreting this pattern as confirmed gene loss.

Previous studies have documented genuine *apxIVA* complexity, including *ISApl1*-mediated disruption, nonsense mutations, fragmented loci and partial *apxIV-S/apxIVS′* duplications associated with mobile elements [15, 20, 21]. *apxIVA* length variation was also recognized in the original description of the toxin, which reported differences in glycine-rich RTX repeat modules among serotypes [54]. Across the 56 complete loci analyzed here, coding sequences ranged from approximately 4.1 to 5.8 kb and recurrent reference-relative gaps were concentrated in the downstream RTX repeat-rich region. In contrast, the experimentally defined SPM and D638–P639 processing boundary were retained in all 56 canonical proteins. The SPM mediates Ca²⁺-dependent autocatalytic processing, and experimental deletion of this region attenuated APP in pigs [17]. Conservation of the SPM across the examined canonical proteins is therefore consistent with functional constraint, although sequence conservation alone does not establish equivalent processing activity in vivo.

The major recurrent length differences occurred downstream of the SPM within the RTX repeat-rich region. RTX repeats coordinate extracellular Ca²⁺, form β-roll structures and can facilitate folding and type I secretion of RTX substrates [55, 56]. Differences in repeat-rich architecture may therefore influence calcium binding, folding, stability, extracellular conformation or antigenic exposure. The extreme C-terminal region containing the predicted type I secretion signal was retained in the natural variants, but secretion efficiency and functional consequences of shorter architectures require experimental testing. Conserved boundaries of the approximately 146-aa and 421-aa reference-relative gaps and their concentration within particular lineages support structured variation, although ancestral-state reconstruction and long-read confirmation would be required to define the underlying evolutionary events.

The additional *apxIVA*-related homolog associated with an IS256-family transposase and a hypothetical protein should be interpreted in the context of previously described *apxIV-S* partial duplications. A transposon-associated *apxIVS′* sequence has been reported in APP [20], so the homolog identified here may represent a related duplication or remodeling event rather than a distinct *apxIVA* family. Its putative SPM-like region was nevertheless highly divergent and lacked the canonical D638–P639 processing boundary. The experimentally demonstrated clip-and-link activity of canonical *apxIVA* therefore cannot be assigned to this homolog from sequence similarity alone. Direct comparison with previously described *apxIV-S/apxIVS′* sequences and functional characterization will be required to resolve its evolutionary and biological significance.

Together, these findings refine interpretation of *apxIVA* as an APP-associated diagnostic locus [15, 20, 57, 58]. Conserved regions remain suitable species markers, whereas the complete gene is more structurally heterogeneous than a single reference sequence suggests. Diagnostic assays should therefore preferentially target regions demonstrated to be conserved across contemporary genomic diversity, and low whole-gene reference coverage should not automatically be interpreted as absence. The functional and antigenic consequences of naturally occurring *apxIVA* length variants remain to be determined.

Several limitations should be considered when interpreting these results. First, the 23 genomic lineages were defined from the core-gene SNP alignment without explicit recombination masking; they should therefore be regarded as operational population clusters, and the contribution of homologous recombination to fine-scale relationships remains to be evaluated. Second, the collection was assembled from diagnostic, surveillance and public-genome sources rather than by population-based sampling, with marked country-level imbalance and epidemiological clustering of 23 L3 isolates from a single Iowa outbreak. Accordingly, lineage frequencies, geographic patterns and adjusted Wallace coefficients describe this collection rather than independent prevalence estimates. Third, structural analysis of *apxIVA* was limited to 56 isolates for which the complete locus, including the full coding sequence and flanking regions, could be assembled on a single contig. This limitation primarily reflects the inherent challenges of short-read sequencing for resolving long and structurally complex genomic regions. The *apxIVA* locus contains repetitive sequences and frequent insertions, which can result in ambiguous mapping and fragmented or incomplete assemblies. Consequently, Illumina short-read sequencing often can not reliably reconstruct the complete *apxIVA* coding sequence together with sufficient upstream and downstream flanking regions. Finally, BLAST-based presence/absence screening cannot fully distinguish genuine gene loss from highly divergent alleles or assembly fragmentation. These constraints do not alter the principal descriptive associations but limit causal, prevalence and functional inference.

Overall, this study places capsule, O-antigen and RTX-toxin variation within a population-genomic framework encompassing 328 APP genomes. Combined K:O genotype showed greater concordance with genomic lineage and ApxI–ApxIII repertoire than K type alone, while *apxIVA*-associated sequence remained broadly conserved but exhibited recurrent lineage-linked structural variation and an additional divergent *apxIVA*-related homolog in some genomes. This finding has important practical implications for diagnostics and surveillance, may inform future vaccine development, and also provides a framework for interpreting virulence-associated genomic diversity and prioritizing future functional studies.

## Author statement

Jiamao Yang: conceptualization, data collection, data curation and analysis, investigation, methodology, project administration, visualization, writing & drafting. Maria Hakimi: methodology, software, data curation and analysis, review & editing. Xiao Hu: data curation and analysis, review & editing. Maria J. Clavijo: review & editing. Marcelo N. Almeida: review & editing. Nubia R. Macedo: data collection, review & editing. Huigang Shen: data collection, investigation, review & editing. Ganwu Li: conceptualization, project administration, supervision, writing, review & editing.

## Acknowledgement

The authors would like to thank the technicians at Veterinary Diagnostic Laboratory at Iowa State University for excellent technical assistance.

## References

1. Hakimi M, Ye F, Saxena A, Hu X, Shen H, et al. Genomic insights into the population structure, antimicrobial resistance, and virulence of Brachyspira hyodysenteriae from diverse geographical regions. Microbiol Spectr;13:e03386–24.

2. Zheng Z, Gorden PJ, Xia X, Zheng Y, Li G. Whole-genome analysis of Klebsiella pneumoniae from bovine mastitis milk in the U.S. Environmental Microbiology 2022;24:1183–1199.

3. Soto Perezchica MM, Guerrero Barrera AL, Avelar Gonzalez FJ, Quezada Tristan T, Macias Marin O. Actinobacillus pleuropneumoniae, surface proteins and virulence: a review. Front Vet Sci;10. Epub ahead of print 30 November 2023. DOI: 10.3389/fvets.2023.1276712.

4. Sassu EL, Bossé JT, Tobias TJ, Gottschalk M, Langford PR, et al. Update on Actinobacillus pleuropneumoniae—knowledge, gaps and challenges. Transboundary and Emerging Diseases 2018;65:72– 90.

5. To H, Frey J, Gottschalk M, Sugiura K, Nagai S. Actinobacillus pleuropneumoniae: An Update on Epidemiology, Biovar, Serotyping, Virulence, and Laboratory Diagnosis. Microbiology and Immunology 2025;69:577–595.

6. Chiers K, De Waele T, Pasmans F, Ducatelle R, Haesebrouck F. Virulence factors of Actinobacillus pleuropneumoniae involved in colonization, persistence and induction of lesions in its porcine host. Vet Res 2010;41:65.

7. Liu F, Peng W, Liu T, Zhao H, Yan K, et al. Biological role of *Actinobacillus pleuropneumoniae* type IV pilus proteins encoded by the *apf* and *pil* operons. Veterinary Microbiology 2018;224:17–22.

8. Baltes N, Hennig-Pauka I, Gerlach G-F. Both transferrin binding proteins are virulence factors in *Actinobacillus pleuropneumoniae* serotype 7 infection. FEMS Microbiology Letters 2002;209:283–287.

9. Shakarji L, Mikael LG, Srikumar R, Kobisch M, Coulton JW, et al. Fhua and HgbA, outer membrane proteins of Actinobacillus pleuropneumoniae: their role as virulence determinants. Can J Microbiol 2006;52:391–396.

10. Nusrat Nahar, Conny Turni, Greg Tram, Patrick J. Blackall, John M. Atack. Actinobacillus pleuropneumoniae: The molecular determinants of virulence and pathogenesis. In: Advances in Microbial Physiology. Elsevier. pp. 179–216.

11. Schuwerk L, Hoeltig D, Waldmann K-H, Valentin-Weigand P, Rohde J. Sero- and apx-typing of German Actinobacillus pleuropneumoniae field isolates from 2010 to 2019 reveals a predominance of serovar 2 with regular apx-profile. Vet Res 2021;52:10.

12. Frey J, Beck M, van den Bosch JF, Segers RP, Nicolet J. Development of an efficient PCR method for toxin typing of Actinobacillus pleuropneumoniae strains. Mol Cell Probes 1995;9:277–282.

13. Kamp EM, Stockhofe-Zurwieden N, van Leengoed LA, Smits MA. Endobronchial inoculation with Apx toxins of Actinobacillus pleuropneumoniae leads to pleuropneumonia in pigs. Infect Immun 1997;65:4350–4354.

14. Frey J, Bosse JT, Chang Y-F, Cullen JM, Fenwick B, et al. Actinobacillus pleuropneumoniae RTX-toxins: uniform designation of haemolysins, cytolysins, pleurotoxin and their genes. Microbiology 1993;139:1723–1728.

15. O’Neill C, Jones SCP, Bossé JT, Watson CM, Williamson SM, et al. Population-based analysis of *Actinobacillus pleuropneumoniae* ApxIVA for use as a DIVA antigen. Vaccine 2010;28:4871–4874.

16. Liu J, Chen X, Tan C, Guo Y, Chen Y, et al. *In vivo* induced RTX toxin ApxIVA is essential for the full virulence of *Actinobacillus pleuropneumoniae*. Veterinary Microbiology 2009;137:282–289.

17. Kuban V, Macek P, Hritz J, Nechvatalova K, Nedbalcova K, et al. Structural Basis of Ca^2+^ - Dependent Self-Processing Activity of Repeat-in-Toxin Proteins. mBio 2020;11:e00226–20.

18. Slivenecka E, Jurnecka D, Holubova J, Stanek O, Brazdilova L, et al. The *Actinobacillus pleuropneumoniae apxIV* operon encodes an antibacterial toxin-immunity pair. Microbiological Research 2025;292:128043.

19. Schaller A, Kuhn R, Kuhnert P, Nicolet J, Anderson T, et al. Characterization of apxlVA, a new RTX determinant of Actinobacillus pleuropneumoniae. Microbiology 1999;145:2105–2116.

20. Srijuntongsiri G, Mhoowai A, Samngamnim S, Assavacheep P, Bossé JT, et al. Novel DNA Markers for Identification of Actinobacillus pleuropneumoniae. Microbiology Spectrum 2022;10:e01311–21.

21. Tegetmeyer HE, Jones SCP, Langford PR, Baltes N. ISApl*1*, a novel insertion element of *Actinobacillus pleuropneumoniae*, prevents ApxIV-based serological detection of serotype 7 strain AP76. Veterinary Microbiology 2008;128:342–353.

22. Bossé JT, Li Y, Fernandez Crespo R, Lacouture S, Gottschalk M, et al. Comparative sequence analysis of the capsular polysaccharide loci of *Actinobacillus pleuropneumoniae* serovars 1–18, and development of two multiplex PCRs for comprehensive capsule typing. Veterinary Microbiology 2018;220:83–89.

23. Beynon LM, Griffith DW, Richards JC, Perry MB. Characterization of the lipopolysaccharide O antigens of Actinobacillus pleuropneumoniae serotypes 9 and 11: antigenic relationships among serotypes 9, 11, and 1. J Bacteriol 1992;174:5324–5331.

24. To H, Maldonado J, Tsutsumi N, Gottschalk M, Frey J, et al. Characterization of Actinobacillus pleuropneumoniae biovar 2 isolates reportedly reacted with the serovar 4 antiserum, and development of a multiplex PCR for O-antigen typing. Veterinary Microbiology 2024;291:110030.

25. Vincent AT, Lacouture S, St-Jean G, Tapia R, Payen S, et al. Atypical Actinobacillus pleuropneumoniae serotype 12 strains with a higher virulence potential. Vet Res 2025;56:149.

26. Xu Z, Chen X, Li L, Li T, Wang S, et al. Comparative Genomic Characterization of Actinobacillus pleuropneumoniae. Journal of Bacteriology 2010;192:5625–5636.

27. Cohen LM, Bossé JT, Stegger M, Li Y, Langford PR, et al. Comparative Genome Sequence Analysis of Actinobacillus pleuropneumoniae Serovar 8 Isolates From Norway, Denmark, and the United Kingdom Indicates Distinct Phylogenetic Lineages and Differences in Distribution of Antimicrobial Resistance Genes. Front Microbiol;12. Epub ahead of print 10 September 2021. DOI: 10.3389/fmicb.2021.729637.

28. Donà V, Ramette A, Perreten V. Comparative genomics of 26 complete circular genomes of 18 different serotypes of Actinobacillus pleuropneumoniae. Microbial Genomics 2022;8:000776.

29. Guitart-Matas J, Gonzalez-Escalona N, Maguire M, Vilaró A, Martinez-Urtaza J, et al. Revealing Genomic Insights of the Unexplored Porcine Pathogen Actinobacillus pleuropneumoniae Using Whole Genome Sequencing. Microbiol Spectr;10:e01185–22.

30. Li S-C, Huang J-F, Hung Y-T, Wu H-H, Wang J-P, et al. In silico capsule locus typing for serovar prediction of Actinobacillus pleuropneumoniae. Microb Genom 2022;8:000780.

31. Angen Ø, Karstensen KT, Vilaró A, Cavaco LM, Langford PR, et al. Serotyping of Actinobacillus pleuropneumoniae based on whole genome sequencing: validation of a bioinformatic tool. Microbial Genomics 2025;11:001434.

32. Ewels P, Magnusson M, Lundin S, Käller M. MultiQC: summarize analysis results for multiple tools and samples in a single report. Bioinformatics 2016;32:3047–3048.

33. Bolger AM, Lohse M, Usadel B. Trimmomatic: a flexible trimmer for Illumina sequence data. Bioinformatics 2014;30:2114–2120.

34. Bankevich A, Nurk S, Antipov D, Gurevich AA, Dvorkin M, et al. SPAdes: a new genome assembly algorithm and its applications to single-cell sequencing. J Comput Biol 2012;19:455–477.

35. Gurevich A, Saveliev V, Vyahhi N, Tesler G. QUAST: quality assessment tool for genome assemblies. Bioinformatics 2013;29:1072–1075.

36. Seemann T. Prokka: rapid prokaryotic genome annotation. Bioinformatics 2014;30:2068–2069.

37. Page AJ, Taylor B, Delaney AJ, Soares J, Seemann T, et al. SNP-sites: rapid efficient extraction of SNPs from multi-FASTA alignments. Microbial Genomics 2016;2:e000056.

38. Nguyen L-T, Schmidt HA, von Haeseler A, Minh BQ. IQ-TREE: A Fast and Effective Stochastic Algorithm for Estimating Maximum-Likelihood Phylogenies. Mol Biol Evol 2015;32:268–274.

39. Kalyaanamoorthy S, Minh BQ, Wong TKF, von Haeseler A, Jermiin LS. ModelFinder: fast model selection for accurate phylogenetic estimates. Nat Methods 2017;14:587–589.

40. Tonkin-Hill G, Lees JA, Bentley SD, Frost SDW, Corander J. Fast hierarchical Bayesian analysis of population structure. Nucleic Acids Res 2019;47:5539–5549.

41. Jansen R, Briaire J, Kamp EM, Gielkens AL, Smits MA. Structural analysis of the Actinobacillus pleuropneumoniae-RTX-toxin I (ApxI) operon. Infect Immun 1993;61:3688–3695.

42. Jansen R, Briaire J, Kamp EM, Smits MA. Comparison of the cytolysin II genetic determinants of Actinobacillus pleuropneumoniae serotypes. Infect Immun 1992;60:630–636.

43. Jansen R, Briaire J, Van Geel AB, Kamp EM, Gielkens AL, et al. Genetic map of the Actinobacillus pleuropneumoniae RTX-toxin (Apx) operons: characterization of the ApxIII operons. Infect Immun 1994;62:4411–4418.

44. Severiano A, Pinto FR, Ramirez M, Carriço JA. Adjusted Wallace Coefficient as a Measure of Congruence between Typing Methods. Journal of Clinical Microbiology 2011;49:3997–4000.

45. Severiano A, Carriço JA, Robinson DA, Ramirez M, Pinto FR. Evaluation of Jackknife and Bootstrap for Defining Confidence Intervals for Pairwise Agreement Measures. PLOS ONE 2011;6:e19539.

46. Jansen R, Briaire J, Kamp EM, Gielkens AL, Smits MA. Structural analysis of the Actinobacillus pleuropneumoniae-RTX-toxin I (ApxI) operon. Infect Immun 1993;61:3688–3695.

47. Bae D-Y, Kang EJ, Cho Y-C, Lim Y, Moon S-H, et al. Actinobacillus pleuropneumoniae in Korea (2015–2025): serovar distribution, toxin gene profiles, antimicrobial resistance, and identification of an apxIICA-deficient serovar 15 profile. J Vet Sci 2026;27:e39.

48. Bossé JT, Li Y, Cohen LM, Stegger M, Angen Ø, et al. Complete genome for Actinobacillus pleuropneumoniae serovar 8 reference strain 405: comparative analysis with draft genomes for different laboratory stock cultures indicates little genetic variation. Microb Genom 2021;7:000687.

49. Vilaró A, Karstensen KT, Cavaco LM, Angen Ø, Solé E, et al. An investigation of the transmission of Actinobacillus pleuropneumoniae within vertically integrated systems using whole genome sequencing. Vet Microbiol 2024;295:110157.

50. Cuccato M, Chiesa F, Mazzone E, Divari S, Cannizzo FT. Characterization of Actinobacillus pleuropneumoniae isolates from pigs in Piedmont, Italy, by whole-genome sequencing: insights into antimicrobial resistance. Front Vet Sci;13. Epub ahead of print 11 August 2026. DOI: 10.3389/fvets.2026.1861912.

51. To H, Tsutsumi N, Kon M, Kawashima N, Koike F, et al. A new subtype of serovar 6, K6b:O3, of *Actinobacillus pleuropneumoniae* based on genotypic analysis. Veterinary Microbiology 2024;298:110291.

52. Komal JPS, Mittal KR. Grouping of *Actinobacillus pleuropneumoniae* strains of serotypes 1 through 12 on the basis of their virulence in mice. Veterinary Microbiology 1990;25:229–240.

53. Nelson MI, Culhane MR, Trovão NS, Patnayak DP, Halpin RA, et al. The emergence and evolution of influenza A (H1α) viruses in swine in Canada and the United States. J Gen Virol 2017;98:2663–2675.

54. Schaller A, Kuhn R, Kuhnert P, Nicolet J, Anderson TJ, et al. Characterization of apxlVA, a new RTX determinant of Actinobacillus pleuropneumoniae. Microbiology 1999;145:2105–2116.

55. Bumba L, Masin J, Macek P, Wald T, Motlova L, et al. Calcium-Driven Folding of RTX Domain β-Rolls Ratchets Translocation of RTX Proteins through Type I Secretion Ducts. Molecular Cell 2016;62:47–62.

56. Baumann U. Structure–Function Relationships of the Repeat Domains of RTX Toxins. Toxins 2019;11:657.

57. Schaller A, Djordjevic SP, Eamens GJ, Forbes WA, Kuhn R, et al. Identification and detection of Actinobacillus pleuropneumoniae by PCR based on the gene apxIVA. Vet Microbiol 2001;79:47–62.

58. Schaller A, Kuhn R, Kuhnert P, Nicolet J, Anderson TJ, et al. Characterization of apxIVA, a new RTX determinant of Actinobacillus pleuropneumoniae. Microbiology (Reading) 1999;145 ( Pt 8):2105– 2116.

